# A benchmark study on current GWAS models in admixed populations

**DOI:** 10.1101/2023.04.27.538299

**Authors:** Zikun Yang, Basilio Cieza Huaman, Dolly Reyes-Dumeyer, Rosa Montesinos, Marcio Soto-Añari, Nilton Custodio, Giuseppe Tosto

**Affiliations:** Taub Institute for Research on Alzheimer’s Disease and the Aging Brain, College of Physicians and Surgeons, Columbia University. 630 West 168 Street, New York, NY 10032, USA; The Gertrude H. Sergievsky Center, College of Physicians and Surgeons, Columbia University. 630 West 168^th^ Street, New York, NY 10032, USA; Department of Neurology, College of Physicians and Surgeons, Columbia University and the New York Presbyterian Hospital. 710 West 168^th^ Street, New York, NY 10032, USA; Unidad de diagnóstico de deterioro cognitivo y prevención de demencia, Instituto Peruano de Neurociencias, Lima, Perú; Instituto de Neurociencia Cognitiva, Arequipa, Perú; Laboratorio de Neurociencia, Universidad Católica San Pablo, Arequipa, Perú

## Abstract

**Objective:** The performances of popular Genome-wide association study (GWAS) models haven’t been examined yet in a consistent manner under the scenario of genetic admixture, which introduces several challenging aspects such as heterogeneity of minor allele frequency (MAF), a wide spectrum of case-control ratio, and varying effect sizes etc.

**Methods:** We generated a cohort of synthetic individuals (N=19,234) that simulates 1) a large sample size; 2) two-way admixture [Native American-European ancestry] and 3) a binary phenotype. We then examined the inflation factors produced by three popular GWAS tools: GMMAT, SAIGE, and Tractor. We also computed power calculations under different MAFs, case-control ratios, and varying ancestry percentages. Then, we employed a cohort of Peruvians (N=249) to further examine the performances of the testing models on 1) real genetic data and 2) small sample sizes. Finally, we validated these findings using an independent Peruvian cohort (N=109) included in 1000 Genome project (1000G).

**Result:** In the synthetic cohort, SAIGE performed better than GMMAT and Tractor in terms of type-I error rate, especially under severe unbalanced case-control ratio. On the contrary, power analysis identified Tractor as the best method to pinpoint ancestry-specific causal variants, but showed decreased power when no adequate heterogeneity of the true effect sizes was simulated between ancestries. The real Peruvian data showed that Tractor is severely affected by small sample sizes, and produced severely inflated statistics, which we replicated in the 1000G Peruvian cohort.

**Discussion:** The current study illustrates the limitations of available GWAS tools under different scenarios of genetic admixture. We urge caution when interpreting results under complex population scenarios.

## Introduction

Genome-wide association studies (GWASs) have successfully identified risk and protective loci in many complex human traits. Among these, binary traits have dominated the pool of explored outcomes, e.g., type 2 diabetes or Alzheimer’s disease (AD). Linear mixed models (LMM), extensively used in GWAS with binary traits, violate the assumption of constant residual variance, leading to inflated type I error. The generalized linear mixed model associated test (GMMAT) ^[1]^ builds logistic mixed models and constructs a score test for the binary traits in GWAS while accounting for population stratification and relatedness via a kinship matrix. Although GMMAT has been shown to be more robust than other LMM approaches with well-controlled type I error rates, it did not address other common limitations, such as imbalanced case-control ratios, a common scenario in the GWAS - especially in population-based studies where affected cases are usually far rarer than controls. Other limitations, such as rare variants, also lead to P-values inflation. To address such limitations, Zhou et al.^[2]^ proposed the Scalable and Accurate Implementation of Generalized mixed model (SAIGE), which includes Saddlepoint approximation (SPA)^[3]^ in the fitting of generalized linear mixed model (GLMM), in order to calibrate the score test accounting for imbalanced case-control ratios and rare variants. Through simulation study and real data analysis, SAIGE shows well-calibrated P values even under these extreme scenarios.

Another pressing issue in GWAS is the under-representation of admixed populations, whose genomes contain segments inherited from multiple ancestral groups. Few GWAS tools have been specifically designed for such complex genetic architecture. Tractor^[4]^, proposed by Atkinson et al., is a scalable framework that incorporates the genetic structure of admixed individuals into large-scale genomics efforts through local ancestry inference, which has been shown to be capable of detecting and modeling ancestry-specific effect sizes. The impact of the local ancestry on association models has also been investigated in a recent publication^[5]^, where the authors compared the performances of Tractor vs. other methods based on the Armitage trend test. However, the latter are fixed-effect models that don’t consider random effects such as the genetic relatedness between individuals. The paper also didn’t account for imbalanced case control ratio, rare variants etc. Therefore, the performances of Tractor have yet to be systematically examined to their full extent.

In general, there is a lack of standardized criterion for benchmarking popular GWAS methods and their results under a variety of key factors, such as minor allele frequency (MAF) heterogeneity, imbalanced case-control ratio, admixture, etc. In this study, we present a benchmark investigation that fills the gap by systematically examining the performances of three popular GWAS models, GMMAT, SAIGE and Tractor, conditional on the factors stated previously. We also applied these tools in an AD study of admixed participants, i.e., Peruvians from the “Genetics of Alzheimer’s disease In Peruvian Populations study” (GAPP) study. Finally, we validated the results using an independent cohort of Peruvians included in the 1000 Genome Project (1000G).

## Methods

### A. Data process

We used a large synthetic dataset using HAPNEST^[6]^, a recently developed software that enabled the generation of a diverse synthetic datasets (using publicly-available reference datasets) of 1,008,000 individuals of 6 ancestry groups. We used the Admixed American (AMR) group from HAPNEST and performed phasing using the 1000 Genome project^[7]^ (1000G) as reference haplotype panel through Shapeit^[8]^ (2.r837). We then used RFMix2^[9]^ (v2.0.3), a discriminative approach that estimates both global and local ancestry using random forests, to inference local ancestry assuming a three-way admixed scenario, i.e., Native-American ancestry (NAA), European (EUR), and African (AFR) through the reference panel Human Genome Diversity project (HGDP)^[10]^. We then filtered out the individuals with significant African global ancestry (i.e., greater than 10%) in order to retain a two-way admixed sample. We again used RFMix2 to estimate local ancestry assuming a two-way admixture of NAA and EUR backgrounds on the remaining 19,234 individuals. We used 19,081 independent genetic variants on Chromosome 20, limiting the analyses to variants with minor allele count (MAC)>10, so that we could investigate the performances of the testing methods to ultra-rare causal variants too. The kinship matrix was computed using PLINK (2.0). The major (i.e., EUR) and minor (NAA) ancestries of the synthetic dataset were modeled opposite to the real Peruvian data (i.e. NAA being the major ancestry and EUR the minor one), to examine a different combinations of admixture.

To apply our methods and perform real-data analyses, we leveraged the GAPP study, a recently established cohort of Peruvian mestizos from Lima and indigenous groups from Southern Peru (Aymaras and Quechuas). Genotyping was conducted on the Infinium Global Screening Array-24 BeadChip, which combines multi-ethnic genome-wide content, curated clinical research variants, and quality control (QC) markers for precision medicine research, extensively detailed in previous publications from our group ^[11]^. We conduct the same procedure by first phasing the genetic data; then, for each individual, global ancestry (NAA, EUR, and AFR) were estimated using the HGDP as the reference panel. After excluding individuals with high African global ancestry, we again inferenced the local ancestry assuming a two-way admixture (NAA and EUR) on the remaining 249 individuals. Variants were filtered with a lower threshold of MAC>5.

We validated the results obtained in the real-data analysis, we used an independent Peruvian cohort from the expanded 1000G^[12]^, again with a two-way admixture profile (NAA and EUR). We filtered out the individuals with high African background and conduct the two-way admixture local ancestry inference (NAA and EUR) as previously discussed. We used 44,847 independent variants on chromosome 1 of the remaining 109 Peruvians using a lower threshold of MAC>10, which represents a more conservative threshold compared to what we implemented in GAPP, ultimately allowing us to investigate the impact of various rare-variant cut-off effect in small sample sized cohorts.

### B. Simulation setting

We conducted a series of simulations to evaluate the performances of the testing methods regarding a variety of different factors within the synthetic cohort from HAPNEST. We evaluated the performances of the testing methods from two perspectives, i.e. the control of type I error rates and the empirical power for detecting the true effect sizes.

1. **Control of type I error**. For testing the control of type I error rates, the binary phenotypes were generated by a logistic mixed model,

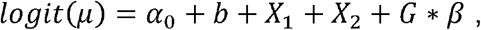

where *G* is the genotype, *β* is the genetic log odds ratio, and *b* is the random effect simulated from a normal distribution N (0, *φ*) with the relatedness matrix *φ*. Two covariates, X_1_ and X_2_, were drawn from Bernoulli (0.5) and standard normal distribution, which represents the discrete and quantitative predictors. The intercept alpha was chosen to represent the corresponding probability of the disease. Under the scenario of the control of type I error, the phenotypes were simulated with *β* = 0. We also simulated three case-control ratios as 1:1, 1:9, and 1:99, denoted as “balance”, “imbalance”, and “extreme-imbalance” scenarios. We then computed and compared the inflation factor *lambda* for each testing method. We also computed the total numbers of the p-values smaller than the genome-wise significance (i.e., 5* 10^−8^) to examine the control of the type I error for each testing method.
2. **Power analysis**. Phenotypes were simulated under the alternative hypothesis, i.e., (3 of the causal variant is not equal to 0. To facilitate the admixture scenario, we simulated that the risk allele was only associated with the NAA ancestry. First, we randomly selected a risk variant conditional on the pre-determined thresholds of MAF. Then, we simulated the phenotype through the probability of disease, which is set to,

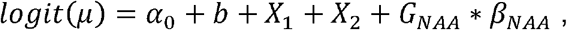

where *G*_*NAA*_ is the genotype matrix associated with the NAA ancestry, X_1_ and X_2_ are the discrete and continuous variables, and b is the random effect. The intercept a_0_ was chosen to reflect the case-control ratios as previously illustrated. We simulated 100 datasets of the corresponding phenotypes for a given MAF and allelic risk effect size of the selected causal variant. The performances of the testing methods were measured through power, which is defined as the proportion of times for which the corresponding test of the causal variant is significant at a given threshold of p values. *MAF*. We categorized the results of the testing causal variants according to the corresponding MAF, such as “ultra-rare” (MAF<0.001), “rare” (0.001<MAF<0.01), “uncommon” (0.01<MAF<0.05), and the common (MAF>0.05). *Varying effect size*. We simulated 100 replicates of simulated genotypes with a logistic model for each allelic effect sizes of the causal ancestry NAA (*β*_*NAA*_) set at 0.25, 0.5, 0.75, 1.0, 1.5, 2.0, 2.5, and 3.0, whereas the effect size of the null ancestry EUR is 0 (*β*_*EUR*_ = 0). *Case-control ratio*. We again consider three case-control ratios: 1:1, 1:9, and 1:99, denoted as “balance”, “imbalance”, and “extreme-imbalance” scenarios.
3. **Impact of heterogeneity of effect sizes between ancestries**. To investigate the impact of the heterogeneity of effect sizes when the causal variants have non-zero causal effect sizes in both ancestries, we conduct a secondary analysis assuming *β*_*EUR*_ = 0.15 and -0.50 < *β*_*NAA*_ < 0.65 increasing by 0.05. The case-control ratio was set at 1:3, and the MAFs of the causal variants of both ancestries were set between 0.1 to 0.2. We measured the testing methods’ performance through power as defined previously in the Power analysis section.
4. **Replication using the Peruvian cohort from 1000G**. To validate the results attained from GAPP, we simulated 100 datasets using Peruvians in 1000G with the same procedure described above and case-control ratio set to 1:3. We examined the total numbers of the variants with p-value smaller than the genome-wide significance.

### C. GWAS methods

For simulations described in 1) and 2), we trained the three GWAS tools, GMMAT, SAIGE, and Tractor. We generated the genomic relatedness matrix (GRM) through PLINK/2.0^[1]^ and provided it to GMMAT, whereas SAIGE creates a sparse relatedness matrix with a default threshold at 0.125 and Tractor does not include the relatedness matrix in its association test. We included the first three principal components to account for population structure. The simulated covariates, *X1* and *X2*, were also provided to the testing models. Tractor fits a logistic regression model including the two ancestry-specific genotypes (while accounting for covariates and local ancestry) then returns the estimated ancestry-specific p-values (in this experiment, we obtained two statistics for EUR and NAA).

For simulations described in 3) and 4), we only trained and compared results from GMMAT and Tractor. In fact, given the case-control ratio being set at 1:3 and the causal variants set as common variants, we did not observe any difference between GMMAT and SAIGE (as expected - data not shown).

For the real-data analysis from GAPP, we again only trained and compared the performances of GMMAT and Tractor, since the case-control ratio was not simulated but derived from real diagnostic status, i.e. 1:3 (58 cases versus 190 controls). We also restricted our analyses to common variants. Therefore, GMMAT and SAIGE produced again overlapping results (data not shown). The first three principal components, age and sex were also used as fixed affects and the GRM as random effect. The local ancestry dosage generated by RFMix was used to implement Tractor as described previously.

## Results

### A. Global ancestries of the testing cohorts in the simulation and real data analysis

**Figure 1**. shows the global ancestry distribution for the three cohorts employed in this project, i.e., the synthetic admixed cohort from HAPNEST, the Peruvians from GAPP, and the Peruvians included in 1000G. The major and minor ancestry of both Peruvian cohorts are NAA and EUR, respectively; the major and minor ancestry of HAPNEST are EUR and NAA, respectively.

**Figure 1.**
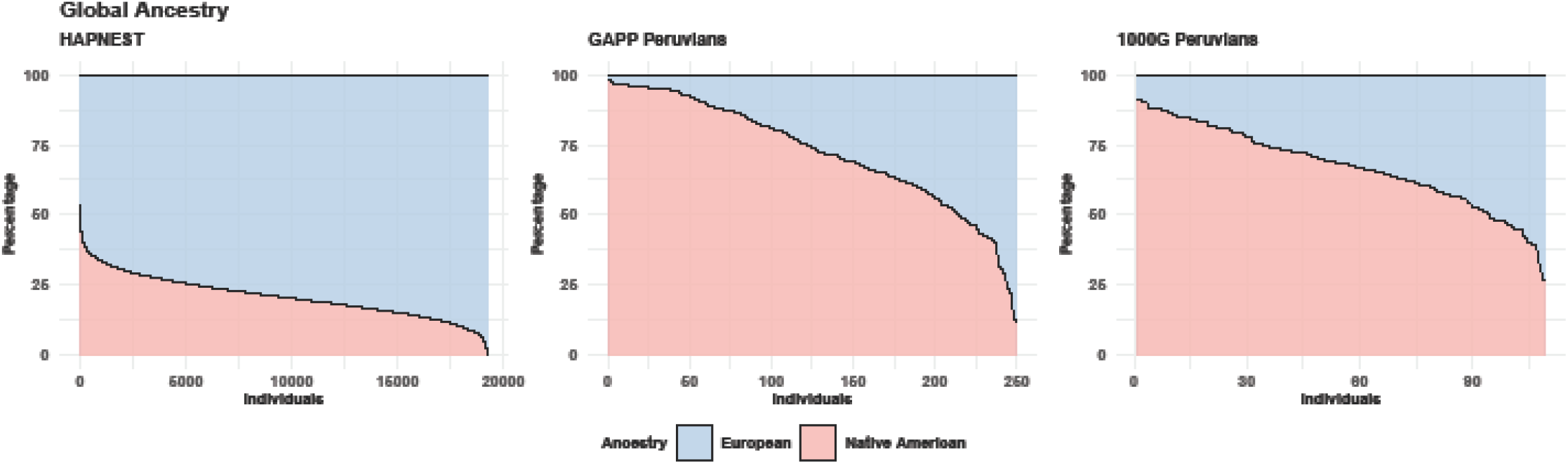
The global ancestries of the individuals included in the HAPNEST, GAPP Peruvian, and 1000G Peruvian.

### B. Type I error rates

Given the large sample size of HAPNEST, all three methods attained acceptable inflation factors with a well-balanced case-control ratio (1:1, **Figure 2**). SAIGE showed well-calibrated inflation factors compared to GMMAT when the case-control ratio shifted to 1:3, whereas Tractor started to show decline in p-values calibration, especially for p-values associated with the minor local ancestry (i.e. NAA). SAIGE and GMMAT both exhibited small inflation in extremely imbalanced case-control ratio (1:99), whereas Tractor showed problematic p-values calibration with inflation factors considerably smaller than 1 for both major and minor global ancestries. As shown in **Table 1**, Tractor produced genome-wide significant results although there were no causal variants being simulated: this ultimately shows that Tractor has high false positive rate (FPR) when the case-control ratio is extremely unbalanced. **Table 1** also reports the median MAFs for the variants associated with false positives results. False positive rates are strongly correlated with low MAFs, with GMMAT’s false positives associated with the ultra-rare variants, whereas Tractor’s false positives extend in the range of rare variants as well. On the other hand, SAIGE was the only method that did not produce false positives under any case-control scenario, and ultimately confirmed the conclusions reached by their authors in its published manuscript.

**Figure 2.**
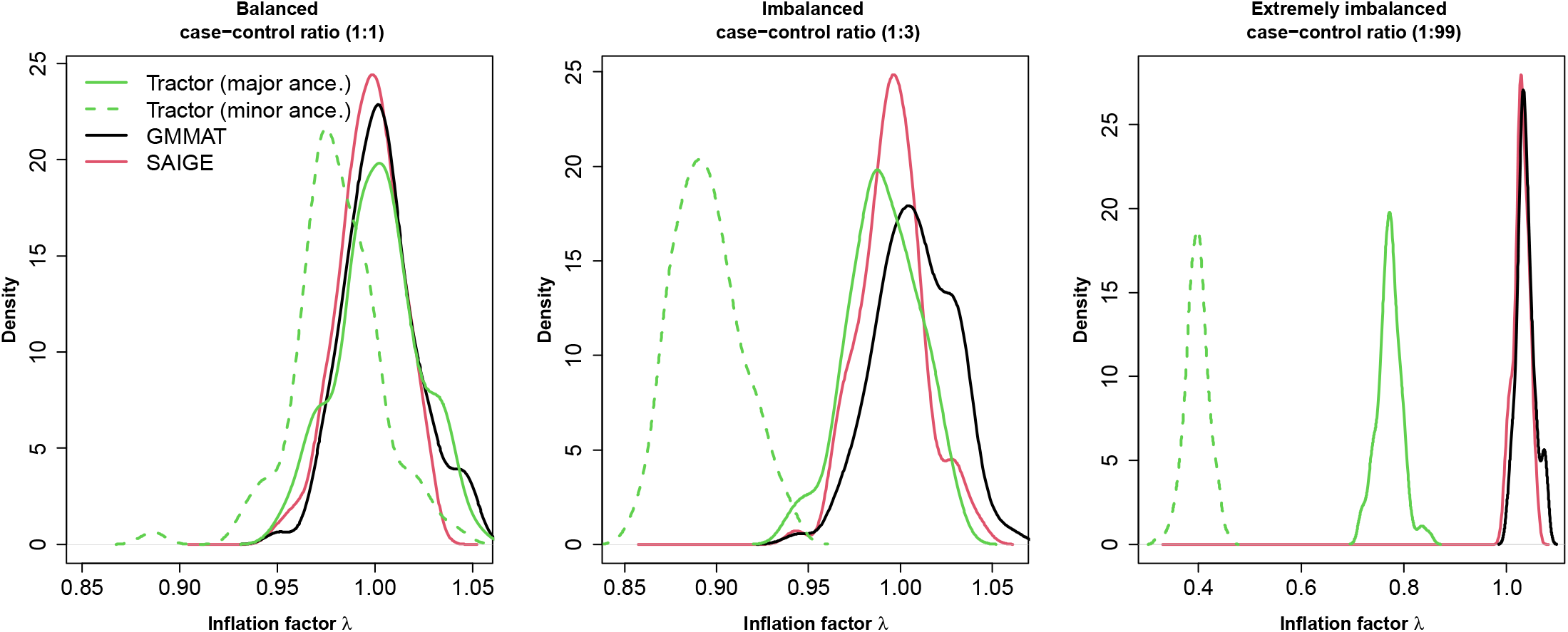
The density function of the inflation factors of the three testing methods over 100 replicates in the simulation scenario of type-I error control.

**Table 1.** The table shows the average number of genome-wide significant variants (“GWV”, p-value ≤5e-8) per 100 replicates, and the median MAF of these variants in the scenario of extremely imbalanced case-control ratio. The MAF of variants identified by Tractor is computed based on local ancestry dosage.

| 100 replicates | GMMAT | SAIGE | Tractor |  |
| --- | --- | --- | --- | --- |
|  |  |  | Major ancestry | Minor ancestry |
| # of GWV | 0.66 | 0 | 1.36 | 3.80 |
| Median MAF | 0.0009 | NA | 0.0034 | 0.0073 |

### C. Power analysis

Under large sample sizes (such as the HAPNEST cohort, **Figure 3**), Tractor showed superior performance in terms of power, i.e., the proportion of causal variants attaining p-values smaller than the genome-wide significance, whereas the performances of GMMAT and SAIGE were virtually similar. For ultra-rare and rare causal variant, Tractor also performed better, although required large true effect sizes. When causal variants were uncommon or common, Tractor again performed better than GMMAT and SAIGE in identifying causal variants with smaller effect sizes under different scenarios of case-control ratio. Tractor also successfully identified the causal ancestry, i.e. the ancestry that the effect sizes are non-zero for simulating the phenotypes.

**Figure 3.**
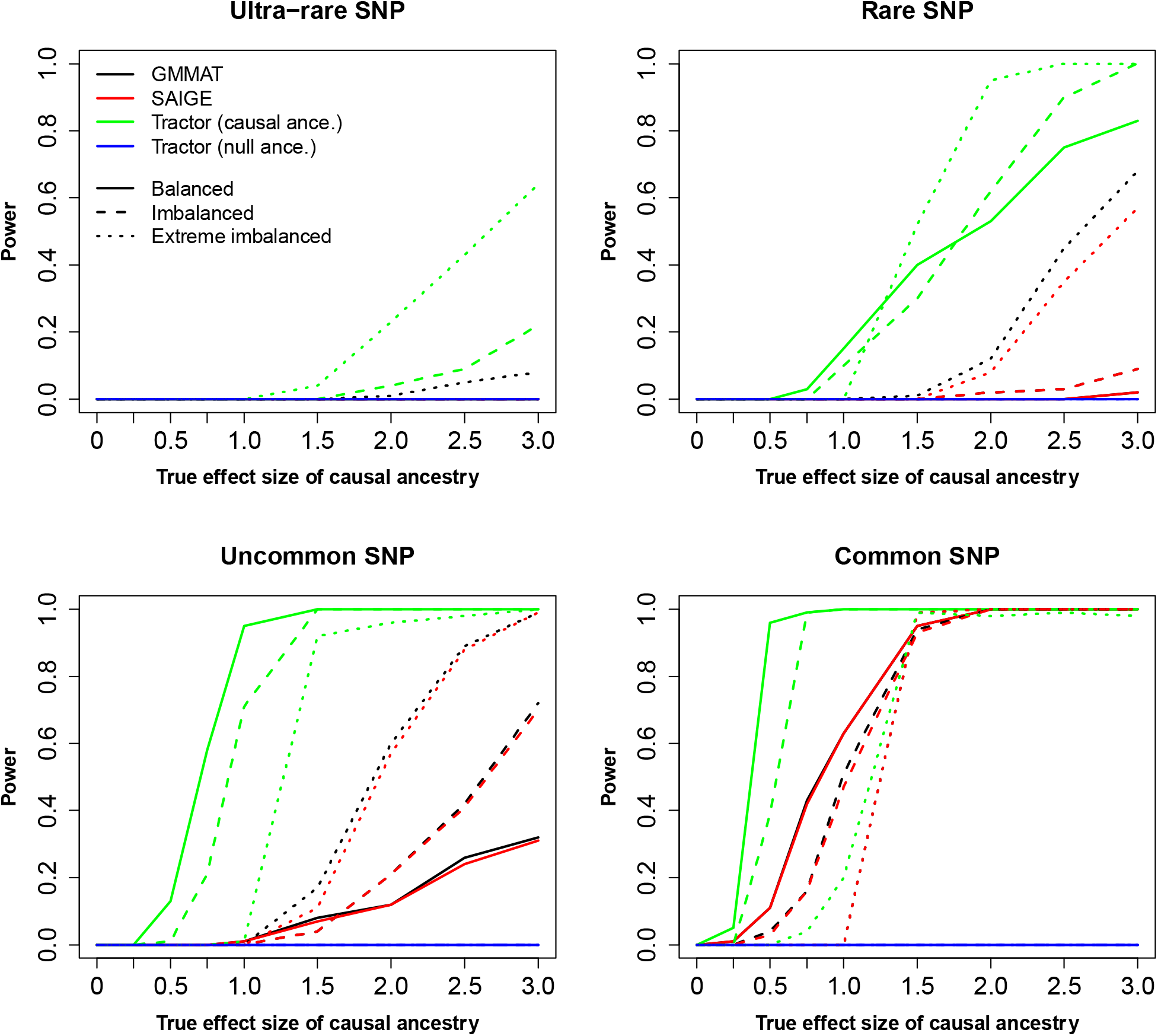
Power calculation of the three methods based on the 19,234 synthetic individuals from HAPNEST. The significance threshold of p-value is set at genome-wide significance (p<5e-8). The causal ancestry, i.e., the corresponding effect size is non-zero, is NAA.

**Table 2** shows that under a balanced case-control ratio (1:1), Tractor controls well FPR as there was no false positive results associated with the null ancestry, i.e. the ancestry that is not associated with the phenotypes when simulating the data. However, when case-control ratio is extremely imbalanced, Tractor retrieved concerningly higher rates of false positives, compared to GMMAT and SAIGE. These false positives were again associated mainly with ultra-rare variants.

**Table 2.** Average number of genome-wide significant variants (“GWV”, p-value ≤5e-8) (and their median MAF) averaging over all 2,400 replicates stratified by case-control ratio. The median MAF for variants identified by Tractor are computed based on local ancestry dosage. We excluded the true causal variants when computing numbers shown in this table.

| 2,400 replicates |  | GMMAT | SAIGE | Tractor |  |
| --- | --- | --- | --- | --- | --- |
|  |  |  |  | Major/Null ancestry | Minor/Causal ancestry |
| Balanced | # GWV | 0.24 | 0.18 | 0 | 0.16 |
|  | Median MAF | 0.033 | 0.033 | NA | 0.14 |
| Imbalanced | # GWV | 0.4 | 0.26 | 0.032 | 0.23 |
|  | Median MAF | 0.04 | 0.034 | 0.00078 | 0.014 |
| Extremely imbalanced | # GWV | 1.18 | 0.2 | 2.03 | 4.93 |
|  | Median MAF | 0.018 | 0.047 | 0.0026 | 0.0062 |

### D. Heterogeneity of effect sizes between ancestry

**Figure 4**. shows that GMMAT is consistently more powerful than Tractor when the effect sizes of the two ancestries are in same direction, i.e., with limited heterogeneity between major (EUR) and minor ancestry (NAA) effect sizes. On the other hand, when the effect sizes of the major and minor ancestry are in opposite directions, i.e., *β*_*EUR*_ = 0.15 while *β*_*NAA*_ < 0, GMMAT is less powerful due to the cancelation of opposite effect sizes. On the contrary, Tractor picks up the causal variants when, for example, the effect size associated with the minor ancestry is large enough to overcome the opposite effect size associated with the major ancestry.

**Figure 4.**
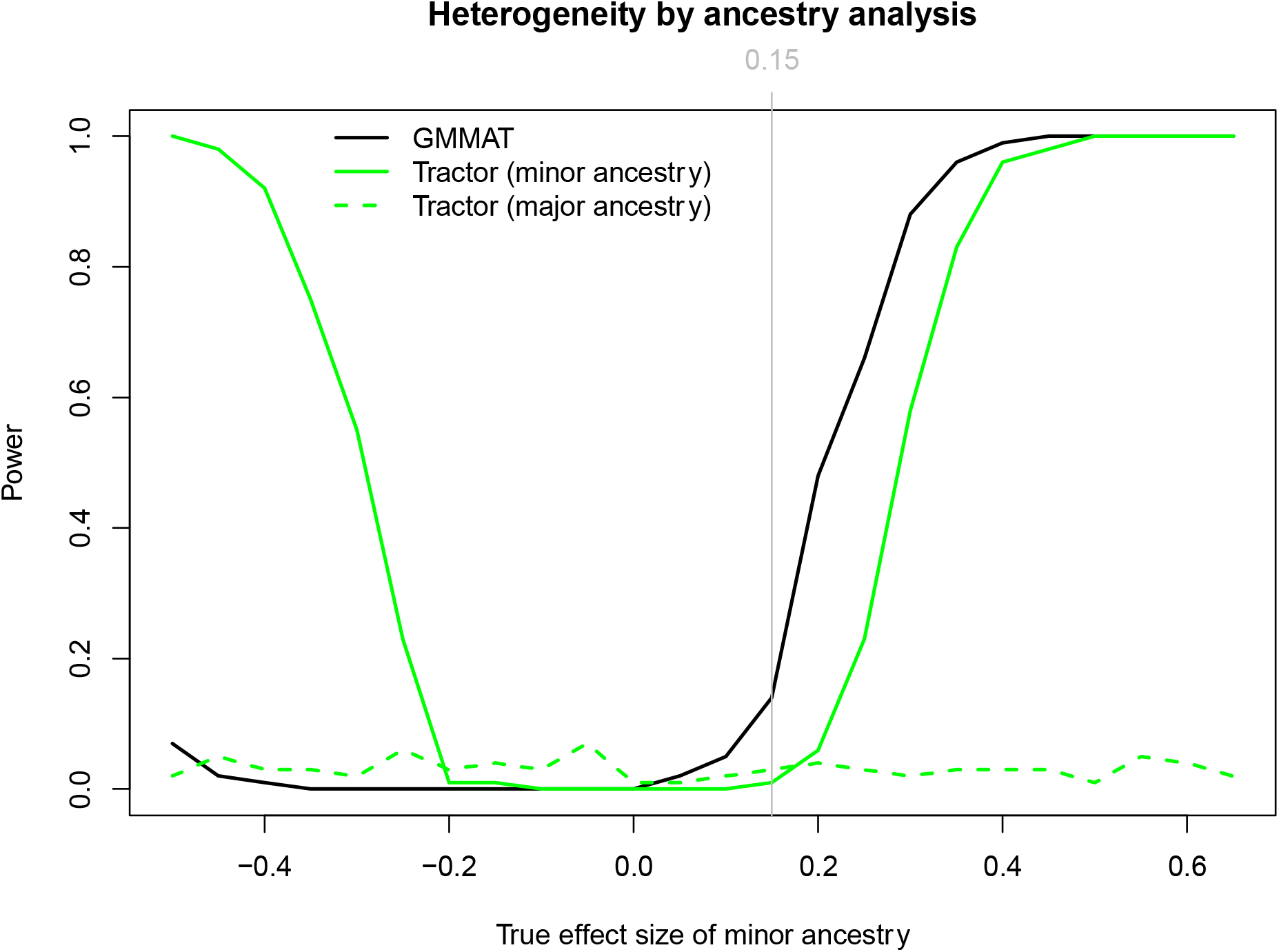
Impact of heterogeneity of the true effect sizes between ancestries on the testing methods. The effect size of the minor ancestry (*β*_*NAA*_) ranges from -0.5 to 0.65 by 0.05, whereas the effect size of the major ancestry (*β*_*EUR*_) is fixed at 0.15.

### E. Real data analysis

**Figure 5a** shows that no genome-wide significance results were achieved in GAPP using GMMAT, likely due to the relatively small sample size (N=249). On the other hand, as shown in **Figure 5b,c** and **Table 3**, Tractor retrieved 31 and 1557 genome-wide significant variants associated with the major (NAA) and minor ancestry (EUR), ultimately showing a severe inflation of p-values and FPR when cohorts have small sample sizes. **Table 3** also shows the correlation between the false positives and MAF.

**Figure 5.**
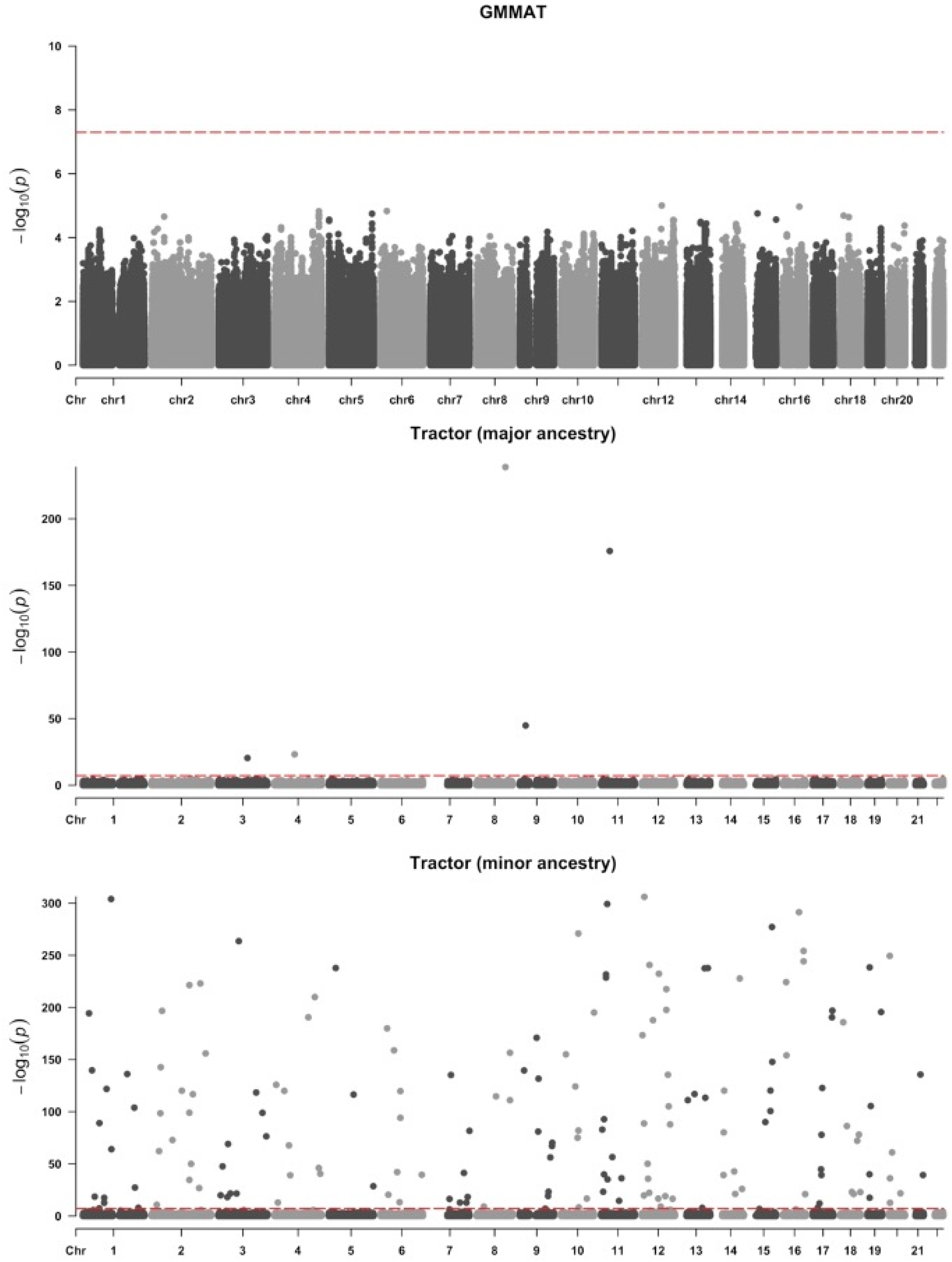
Manhattan plots of AD GWAS produced by GMMAT (**a**) and Tractor (**b**: NAA and **c**: EUR) in the GAPP cohort. The total number of tested variants is 4,492,989.

**Table 3.** Total numbers of genome-wide significant variants (“GWV”, p-value ≤5e-8) identified by GMMAT and Tractor in GAPP.

|  | GMMAT | Tractor |  |
| --- | --- | --- | --- |
|  |  | Major ancestry | Minor ancestry |
| # GWV | 0 | 31 | 1557 |
| Median MAF | NA | 0.031 | 0.014 |

We replicated conclusion about Tractor’s false positives by leveraging the Peruvians from 1000G (N=109). Here, Tractor again returned worrisome results, i.e., 30 and 4,031 genome-wide significant variants for major (NAA) and minor ancestry (EUR) respectively, while no such variants were identified by GMMAT. **Table 4** shows that, likely due to small sample size, the deconvolution of the genotypes by local ancestries creates not only rare variants but also monomorphic variants associated with the minor ancestry, even though all variants were filtered by MAF > 0.01 in the overall sample.

**Table 4.** Total numbers of genome-wide significant variants (“GWV”, p-value ≤5e-8) identified by GMMAT and Tractor in 1000G.

|  | GMMAT | Tractor |  |
| --- | --- | --- | --- |
|  |  | Major ancestry | Minor ancestry |
| # GWV | 0 | 30 | 4031 |
| Median MAF | NA | 0.056 | 0 |

## Discussion

In this study we employed large and small cohorts of synthetic and real-data admixed individuals, and benchmarked the results obtained by three popular GWAS methods, GMMAT, SAIGE, and Tractor under various scenarios.

When power was investigated, we observed optimal performance by Tractor in identifying and modeling the causal effect sizes, compared to the GMMAT and SAIGE. The superiority of Tractor was particularly evident when large heterogeneity existed in terms of effect sizes between ancestries. Our conclusion is in line with results reported in the original paper^[4]^ and the other benchmark paper comparing Tractor to the Armitage trend test^[5]^. However, the deconvolution of the genotype matrix is analogous to reducing the sample size, which ultimately leads to inferior performances of Tractor, especially when the heterogeneity across effect sizes is not large enough (Figure 4).

When studying the type I error control, SAIGE generated well calibrated p-values even under extreme situations, such as rare variants and imbalanced case-control ratios. GMMAT showed small inflation of p-values and produced false positives only under extreme scenarios. Lastly, the calibrations of p-values by Tractor, especially for the minor ancestry, were greatly affected by the imbalanced case-control ratios, even when sample sizes are large - as is the case of the HAPNEST cohort.

Given HAPNEST large sample size and balanced case-control ratio, Tractor shows well-controlled FPR, but produced substantial false positive results associated with the rare variants and extreme case-control ratio in both simulation studies (type I error control and power analysis). It must be emphasized that the simulation study of type I error control in HAPNEST included only 19,081 variants from chromosome 1 leading to an average of 5.16 false positives across major and minor ancestries reported by Tractor. Hence, if Tractor was used for whole-genome association tests, we could expect thousands or tens of thousands of false positives. The mishandling of rare variants is further aggravated when the sample size is small, such as the case of real-data analysis (GAPP and 1000G cohorts), because the deconvolution of the genotype matrix further creates rare variants in the local ancestry dosages. In fact, in an extreme scenario all minor alleles can be attributed to one ancestry, which leaves the other ancestry with monomorphic variants only, ultimately producing outstandingly false positive results. Further, it should be noted that Tractor may not be suitable for analyzing related samples, as Tractor does not account for kinship, which could lead to inflated p-values or false positives.

In summary, we acknowledge the improvement achieved by Tractor in identifying ancestry-related causal variants, by leveraging the unique genetic structure of admixed populations. However, we want to caution the usage of Tractor under extreme circumstances, especially in small sample sizes and when the deconvolution of genotype matrix introduces additional issues in terms of allele frequency. This study demonstrates the importance of considering imbalanced case-control ratio, rare variants and sample size, and ultimately addresses the major challenges for the development of future GWAS methods in admixed populations.

## Acknowledgments

The National Institutes of Health (NIH), National Institute on Aging (NIH-NIA) supported this work through the following grants: R56AG069118, R56AG066889, R56AG059756, U19AG074865, R01AG082009.

## Author contributions

ZY, GT contributed to the conception and design of the study.

GT, NC, ZY, BH, DRD, RM, MSA contributed to the acquisition and analysis of data.

GT, ZY contributed to drafting the text or preparing the figures.

